# Consequences of pharmacophagous uptake from plants and conspecifics in a sawfly elucidated using chemical and molecular techniques

**DOI:** 10.1101/2021.02.09.430406

**Authors:** Sarah Catherine Paul, Alice B. Dennis, Lisa Johanna Tewes, Jeanne Friedrichs, Caroline Müller

## Abstract

Pharmacophagy involves the sequestration of specialised plant metabolites for non-nutritive purposes and commonly occurs in insects. Here we investigate pharmacophagy in the turnip sawfly, *Athalia rosae*, where adults not only collect specialised metabolites (clerodanoids) from a plant (*Ajuga reptans*), but also from the exterior of conspecifics via fighting. Using behavioural assays, chemical analytics, and RNAseq we show that when individuals nibble on conspecifics that have already acquired clerodanoids from *A. reptans* leaves, this nibbling results in the transfer of compounds between individuals. Furthermore, unlike other pharmacophagous insects, the acquisition of clerodanoids by *A. rosae* from the leaves of *A. reptans* does not induce the upregulation of known detoxification or sequestration genes and pathways. In contrast, pharmacophagous nibbling on conspecifics results in the upregulation of metabolic pathways associated with elevated metabolic rates and increased energy consumption. It therefore seems that individuals attack conspecifics to acquire clerodanoids despite the apparent metabolic costs of this form of pharmacophagy compared to clerodanoid uptake from a plant. Changes in the metabolic phenotype of *A*.*rosae* individuals consequently has profound consequences for social interactions with possible ramifications for their social niche.

**Summary statement:** The turnip sawfly (*Athalia rosae*) gains potentially beneficial compounds from the leaf surface of non-food plants (e.g. *Ajuga reptans*), but can also steal these compounds from conspecifics via aggressive nibbling.

## Introduction

Competition between individuals over resources such as food (Jakob, 1994) or mates (Karino et al., 2005) is well known, but comparatively less explored is competition between individuals over chemical compounds, for example those acquired from plants for non-nutritional purposes, a behaviour known as pharmacophagy (Boppré, 1984). Pharmacophagously collected phytochemicals, usually specialised plant metabolites, can have significant fitness benefits for individuals, aiding in antipredator defence (Rojas et al., 2017), resistance to pathogens and parasites (Tan et al., 2019), and successful mating (Iyengar et al., 2001). Consequently, sources of relevant phytochemicals may become an arena for contest and combat, especially when scarce or patchily distributed throughout an individual’s environment. Such contests are likely to be especially common in insects which are known to operate in a highly chemically mediated world (Tabata, 2018), and whose pharmacophagous behaviour is some of the best documented in the animal kingdom (Heckel, 2014). By taking up plant compounds and potentially modifying them, individuals change their chemical phenotype with potential ramifications for interactions with conspecifics and therefore their social niche (Müller et al., 2020). Here we investigate contest interactions over a resource acquired by pharmacophagy and look at how prior sequestration of plant compounds influences motivation to fight, whilst also assessing the physiological consequences of both pharmacophagy and fighting behaviour.

Following predictions from game theory, the occurrence and outcome of contest interactions between two conspecifics (in this case over phytochemical resources) depends on the balance between the costs and benefits of engaging in a fight for each individual (Enquist and Leimar, 1987; Maynard-Smith and Price, 1973). This balance in turn is closely linked to the likelihood of each individual winning a fight, its so-called resource holding potential (RHP; Arnott and Elwood, 2009), and the value of the contested resource to each competitor (resource value or RV; Arnott and Elwood, 2008). RHP is commonly determined by the size of an individual and its weapons or factors linked to its condition, such as persistence capacity or strength (for review see Vieira and Peixoto, 2013). For example, larger competitors can be more likely to win an interaction against smaller or less experienced individuals (Chamorro-Florescano et al., 2011). However, RV can influence contest occurrence and outcome (Stockermans and Hardy, 2013); for example, females with offspring are more likely to win contests against males over burrows than females without offspring (Figler et al., 1995). One might therefore predict that where differences in RHP are minimal, when competing over phytochemical resources, including those that confer chemical defence, the individual most likely to win the interaction would be the one that currently has no or low levels of chemical defence. That is the costs of fighting, increased energetic expenditure (Hack, 1997) and predation risk (Jakobsson et al., 1995), will only be outweighed by its benefits for those individuals that will accrue possible fitness gains, e.g. via defence against predators or parasitoids.

Pharmacophagy itself can carry costs, particularly when it comes to the sequestration of the more toxic specialised plant metabolites (reviewed in Ruxton et al., 2019). Although the extent to which sequestration is actually costly varies considerably between species (Zvereva and Kozlov, 2016), there are some clear cases where dealing with high toxin load during pharmacophagy incurs a physiological burden (Camara, 1997; Züst et al., 2018). The variability between species does however suggest that sequestration costs likely depend heavily on the nature of the compound/s being obtained and on an organism’s physiological mechanisms for dealing with them (Heckel, 2014). In terms of the mechanisms involved in sequestration, a few detoxification and sequestration pathways have been identified, involving the expression of genes from a number of common groups including cytochrome P459 monooxygenases (P450s), glutathione S-transferases (GSTs), UDP-glucuronosyltransferases (UDP), and ABC transporters (Erb and Robert, 2015; Vogel et al., 2014). An understanding of the mechanisms underlying such sequestration costs in contest scenarios is crucial as it may inform an individual’s motivation to fight (RV) over pharmacophagously acquired resources. For example, an individual that has already obtained some defence chemicals may still require access to a resource if the defence is concentration-dependent (Burdfield-Steel et al., 2019), meaning that its RV remains high despite previous access. In contrast, an individual who has collected larger amounts of compounds by pharmacophagy may have reached a threshold of either sequestration costs or an adequate level of defence.

Adults of *Athalia rosae* (Hymenoptera: Tenthredinidae) pharmacophagously acquire neo-clerodane diterpene compounds, from now on called ‘clerodanoids’, from non-food plants such as *Ajuga reptans* (Lamiaceae) (Nishida et al., 2004). Ingestion of these compounds changes their metabolic phenotype, i.e. metabotype, and has positive effects on both female mating success and predator deterrence (Amano et al., 1999; Nishida and Fukami, 1990). Like many other pharmacophagous species, *A. rosae* adults also seem to alter some of the compounds that they take up (Nishida et al., 2004). However, what makes them unique is that in addition to nibbling on leaves adults attack and nibble on conspecifics in what seems to be an attempt to gain these compounds from the body of their opponent. Here we aim to formally document this agonistic nibbling behaviour and to test whether: 1) it is stimulated by a conspecific opponent’s acquisition of clerodanoids and if it varies with an individual’s own clerodanoid status, 2) results in the transfer of compounds between individuals, and 3) leads to differential gene expression between individuals that collect clerodanoids from leaves or from conspecifics. Specifically we predict that 1) the strongest determinant of successful nibbling will be disparity between individuals in clerodanoid acquisition, i.e. nibbling will be highest for individuals lacking clerodanoids facing individuals that already had access, 2) nibbling will result in the transfer of compounds between individuals, and 3) pharmacophagy on leaves will result in the upregulation of canonical detoxification and sequestration pathways and genes, whereas pharmacophagy on conspecifics will induce the upregulation of metabolic pathways related to increased energy production and stress.

## Methods

### Rearing scheme of *A. rosae*

Adults of *A. rosae* (F0) were collected in a meadow in Verl, Germany (51°52’23.0”N 8°33’32.0”E: July 2018) and used as experimental individuals. These adults (N = 60) were kept in two groups (A and B), provided with *Sinapis alba* plants for oviposition and *Brassica rapa* (var. *pekinensis*) plants as food for emerging F1 larvae (both Brassicaceae). All F1 larvae were transferred to individual pots containing ≃ 30 g sterilised soil for pupation. Post emergence, F1 individuals were mated with an individual of the opposite breeding group (e.g. F1♀A × F1♂B). Mated (N = 30) and virgin (N = 30) F1 females were provided with a *S. alba* plant for oviposition and a honey-water mixture as food (1:50). F2 larvae were placed in ventilated containers (25 cm × 15 cm × 10 cm), and provided *ad libitum* with middle-aged *B. rapa* leaves and moistened tissue. As for F1 larvae, F2 larvae were transferred into soil pots for pupation and after emergence F2 adults were placed in individual Petri dishes (60 mm × 15 mm) with honey water-infused tissue paper as a food supply. Adults were kept at ~5 °C until commencement of contest bioassays. Rearing took place in a climate chamber (temp: 20 °C:16 °C (16 h:8 h), light:dark (16 h:8 h), 70 % r.h.). Host plants were grown from seed in a greenhouse (no climate control, light:dark 16 h:8 h).

#### 1) Contest bioassays

Contest bioassays were carried out between male *A. rosae* with pairs established across four different treatment levels with differing focal and opponent clerodanoid acquisition status (i.e. with clerodanoids (C+) or without clerodanoids (C−); Fig. 1). Two days prior to the start of the trials F2 adult *A. rosae* (3-14 days post eclosion) were removed from the refrigerator, weighed to the nearest 0.01 mg (Sartorius AZ64, M-POWER Series Analytical Balance), provided with a honey–water mixture, and individuals in the C+ treatment were additionally provided with a small piece (1 cm^2^) of a leaf of *A. reptans* (Lamiaceae). Plants of this species had been collected close to a forest (52°01’58.2”N 8°29’04.5”E) and were kept outdoors. Males were marked (post-weighing and after being briefly chilled at 5 °C) at the top of the thorax and anterior to the wing joints with a small colour spot of quickly drying coloured lacquer (Maybelline Colour Show 60 Seconds, Maybelline, New York) in either pink (16R 200) or blue (16R 400). These lacquers were chosen for similarity in chemical composition and colour luminance, but difference in chromatic components, and each colour was distributed equally among treatments. Individuals from each clerodanoid acquisition status (i.e. C+ or C−) were handled with different forceps and paired individuals were matched for size (with mass as a proxy for size). In addition, sib-sib pairings were avoided and each pair was assigned to one of the four treatment levels in a way that maximised the even spread of individuals from different parents across treatments. Contests began when individuals were added to opposite sides of the experimental arena (Petri dish) and were filmed for 25 min using a Sony HDR-CX410VE Camcorder (AVCHD −1920 × 1080 −30 fps). The software BORIS v. 7.9.7 (Friard and Gamba, 2016) was used to analyse the video data, which was done blind by a single observer. Successful nibbling behaviour was classified as the mouthparts of one individual making contact with the body of another individual and its occurrence and duration were recorded at 0.3x the original speed and at 2x display magnification to ensure correct timing and categorisation. At the end of each trial individuals were isolated in Petri dishes, provided with fresh honey–water mixture and kept for another 48 h, to allow for the metabolism of any clerodanoids gained through nibbling. Afterwards they were frozen at −80 °C until chemical analysis (see below).

**Figure 1.**
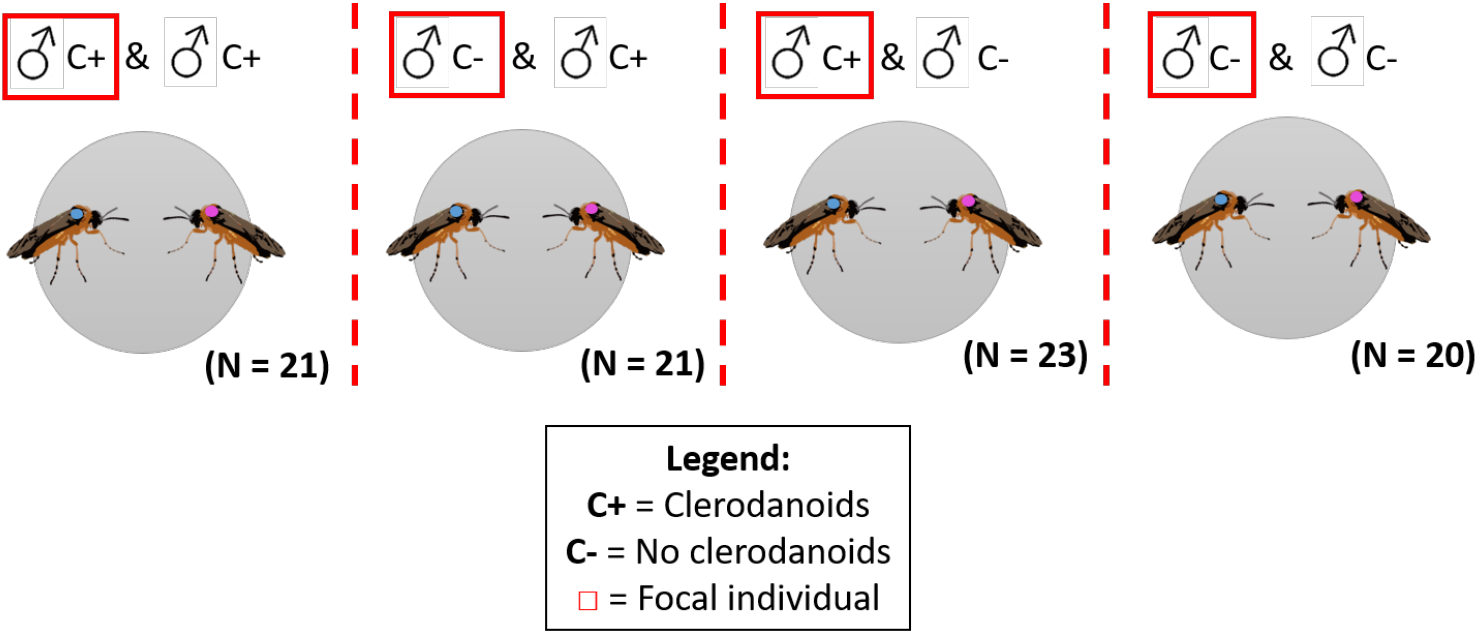
Different treatments and number of pairs used (N) in the male-male contest assays with four treatment levels, treatment levels outlined at the top and the focal individual in each contest highlighted using a red box for further clarification. C+ individuals (with clerodanoids) were provided with a small piece (1 cm^2^) of *Ajuga reptans* leaf material 48 hours prior to the assays, whereas C− individuals (no clerodanoids) were not. Although not shown in the diagram above, within each level/combat pair (e.g. C+ *vs* C−) half of the trials were carried out with the focal individuals marked with pink lacquer (e.g. C+ pink and C− blue) and half with the focal individuals marked with blue (e.g. C+ blue and C− pink), to prevent the potential confounding effects of the lacquer colour.

#### 2) Analysis of pharmacophagy and conspecific transfer of clerodanoids

To test whether nibbling on *A. reptans* leaves resulted in the acquisition of clerodanoids by *A. rosae*, C+ (N=8) and C−(N=7) individuals from the male-male contest trials (C+C− and C−C− treatments respectively), which had not engaged in any conspecific nibbling behaviour, were chemically analysed. To test whether conspecific nibbling resulted in the acquisition of clerodanoids, C− individuals that had nibbled on C+ individuals (AC+ individuals, N=13) were compared to C− individuals that had either not nibbled on C+ individuals (C+C− or C−C+ treatment, N=3) or had nibbled on a C− individual (C−C− treatment, N=7). To extract clerodanoids from *A. rosae* individuals, whole insects were frozen, lyophilised (for 48h) and homogenised. Individual samples were extracted in 130 µl ethyl acetate (LC-MS grade, VWR, Leuven, Belgium) for 10 min, centrifuged and 120 µl of the extract dried under reduced pressure (15 min at 30 °C). Dried extracts were resuspended in 75 µl 90% methanol LC-MS grade, Fisher Scientific, Loughborough, UK) and filtered with syringe filters (polytetrafluoroethylene membrane, 0.2 µm pore size, Phenomenex, Torrance, CA, USA). Lyophilised samples of *A. reptans* leaf material were prepared likewise to identify potential clerodanoids taken up by the insects. All samples were separated on a Kinetex XB-C18 column (1.7 µm, 150 × 2.1 mm, with guard column, Phenomenex) and analysed using an ultra high performance liquid chromatograph (Dionex UltiMate 3000, Thermo Fisher Scientific, San José, CA, USA) coupled to a quadrupole time of flight mass spectrometer (compact, Bruker Daltonics, Bremen, Germany) used with negative electrospray ionisation (UHPLC-QTOF-MS/MS, see S1 for settings). Chromatograms were processed using the software Compass Data Analysis 4.4 (Bruker Daltonics). Candidate compounds which may have been taken up (and modified) by the insects from the plants were identified firstly by comparing the chromatograms of C+ and C− individuals, as the signals were likely to be stronger in C+ than AC+ individuals, and then by comparing *A. reptans* leaf extracts to the extracts of all *A. rosae* individuals. The peak areas of two putative clerodanoids were then manually integrated from the extracted ion chromatograms by searching for the respective m/z within a range of ± 0.02 m/z.

#### 3) Gene expression

##### RNAseq: sample collection, library preparation, sequencing, and alignment

A total of 18 adult males (4-10 days old), comprising six biological replicates per treatment level, were taken for transcriptome sequencing to assess the effects of the two different types of clerodanoid exposure, nibbling on an *A. reptans* leaf (C+) and nibbling on a conspecific (AC+), in comparison to the control (C−). C+ individuals were exposed to a fresh *A. reptans* leaf in a clean Petri dish lined with damp filter paper to prevent desiccation, and 60 min after they commenced feeding on *A. reptans* leaves individuals were frozen in liquid nitrogen. C− individuals were transferred to clean Petri dishes lined with damp filter paper and were then frozen in liquid nitrogen approximately 60 min later. To produce AC+ individuals, C− individuals were set up in a contest against C+ individuals that were prepared using the same method as for chemical analysis (methods above). C+ individuals used in the interactions were chosen so that they would be smaller than C− individuals, and a small section of wing in the C+ individual was clipped so that C− and C+ individuals could be easily distinguished. One hour after C− individuals first nibbled on C+ individuals in the contest (making them AC+), they were frozen in liquid nitrogen. Individuals were chosen in a way that maximised the equal spread of siblings across treatments. All samples were stored at −80 °C until extraction approx. 1 month later.

RNA was extracted from whole individuals using an Invitrogen PureLink ™ RNA Mini Kit (ThermoFisher Scientific, Germany) with a DNase step (innuPREP DNase I Kit, analyticJena, Jena, Germany) and diluted to 25 µl using RNAse free water to a concentration of ~30-100 ng/µl per sample. RNA quality was measured using both a bioanalyzer 2100 (Agilent, CA, United States) and Xpose (PLT SCIENTIFIC SDN. BHD. Malaysia). Novogene (Cambridge, UK) carried out library preparation (with a Ribo-Zero for rRNA removal) and sequencing using a NovaSeq6000 and S4 Flowcells (Illumina, CA, United States). Sequencing of the 18 adults across 3 different treatments yielded a total of 1.24 billion reads with an average of 68 million reads per sample (SEM: ±2million reads) and an average GC content of 40%. The quality of the sequenced libraries was assessed by FastQC software version 0.11.9 (Andrews, 2010) run both before and after cleaning the raw reads. Illumina specific adapter sequences, sequences shorter than 75 nucleotides, and poor-quality sequences (quality <4 over a 25 bp sliding window) were removed using Trimmomatic (Bolger et al., 2014). Reads retained after cleaning were mapped to the annotated *A. rosae* genome, AROS v.2.0 (GCA_000344095.2), with RSEM v1.3.1 (Li and Dewey, 2011); mapping within this was performed with STAR v2.7.1a (Dobin et al., 2013). Expected counts from STAR were used for differential expression analysis.

Unmapped reads were also extracted to check for differentially expressed genes not present in the reference genome. Unmapped reads were extracted from the RSEM-produced BAM file using samtools view (Li et al., 2009) and converted to fastq (using bamtools bamtofastq; Barnett et al., 2011). A total of 247,997,896 reads were extracted and *de novo* assembled using Trinity (default settings) (Grabherr et al., 2011). The unmapped reads were mapped back to this reference and expected read counts extracted using eXpress (Roberts et al., 2011). Transcripts with >10,000 mapped reads were identified using BLASTN (default settings, limited to one match per gene and 1e-1) against the NCBI nucleotide database. As with mapped reads a DE analysis was also carried out for unmapped reads.

##### Differential expression and gene set enrichment analysis

Differential expression analysis and the visualisation of results were carried out in R 4.0.2 (2020-06-22) --“Taking Off Again” using DESeq2 version 1.28.1 (Love et al., 2014). Reads were imported from the RSEM output to DESeq2 using Tximport version 1.16.1 (Soneson et al., 2015) to account for read length and summarise transcript-level mapping for analysis at the gene-level. Prior to differential expression analysis, genes with zero counts in all samples and those with low counts (<10) in less than a quarter of samples (= 6 samples) were dropped from the dataset. Principle component analyses (PCA) and heatmaps of both the vst (variance stabilising transformation) and rlog transformed data were used to scan for any outlying samples, and one sample (AC+ sample 18) was dropped from the analysis based on its large overall expression differences relative to all other samples (Gierliński et al., 2015). DESeq2 allows for both the normalisation of read counts (accounting for differences in library depth/sequencing depth, gene length, and RNA composition) and the shrinkage of gene-wise dispersion (accounting for differences in dispersion estimates between genes) whilst modelling a negative binomial generalised linear model (glm) (to correct for overdispersion) of predictors for each gene (Love et al., 2014). The predictor variable was modelled as ‘treatment’ with the three different levels of clerodanoid exposure (AC+; C+; C−). We made all three of the individual pairwise comparisons within the model built from all samples (Fig. 4). To test whether the differential expression of a gene between two treatment levels was significantly different to zero, pairwise contrasts were performed using a Wald Test and shrunken LFC values with apeglm version 1.10.1 (Zhu et al., 2019). P-values were adjusted for multiple testing via the Benjamini and Hochberg procedure using a false discovery rate (FDR) of 0.05 (Benjamini and Hochberg, 1995). Significantly differentially expressed genes (both relatively up-or down-regulated) were then extracted and filtered with a p-adjust of <0.05. The relationship between the differentially expressed genes for each pairwise comparison was visualised using a Venn Diagram (package=Venn.Diagram, version 1.6.20). The expression of the significantly differentially expressed genes from each comparison was visualised in a heatmap using normalised counts scaled per gene (scale=“row”) (pheatmap, version 1.0.12).

To characterise the differential expression at the pathway level, KEGG terms were allocated to the gene expression data using the KEGG Automatic Annotation Server (KAAS) (KAAS; Moriya et al., 2007). A bi-directional best hit KAAS was run using the predicted genes from the *A. rosae* genome, with 40 different insect species selected for reference (S2). 65% of the annotated genes for which we had read counts were assigned at least one KEGG term. As multiple genes matched to the same KEGG term, the normalised reads were summed across all genes that matched to each term. Furthermore, in the small number of cases (<30), in which multiple KEGG terms were assigned to the same gene, the read counts were duplicated for each individual KEGG term. A gene set enrichment analysis (GSEA) was then carried out on the normalised counts using GAGE (Luo et al., 2013) and pathview (Luo and Brouwer, 2013) in R using an FDR-adjusted p-value cut-off of <0.05 to identify differentially expressed pathways.

### Statistical analysis

All data were analysed using R version 3.6.3 (R Core Team, 2020). Alpha level was set at 0.05 for all tests and model residuals were checked for normality and variance homogeneity. Stepwise backwards deletion using Chi-squared ratio tests (package: MASS, version 7.3-51.6) was employed to reach the minimum adequate model (Crawley, 2012). Posthoc analyses were carried out using the package ‘multcomp’ (version 1.4-13, Hothorn et al., 2008).

### Contest bioassays

Differences in the occurrence (number of events) and duration of successful nibbling were assessed at both the treatment or contest level (C+ vs C+; C+ vs C−; C− vs C−) and the level of the individual (focal individual = C+/C−, non-focal individual = C+/C−; following guidance in Briffa et al., 2013). Variation in the occurrence of successful nibbling between the different treatment levels was tested using a negative binomial hurdle model to account for zero inflation (dist=glm.nb, link=logit, package: DHARMa, version 0.3.2.0, Hartig, 2020), where nibbling occurrence (count) was the response and treatment the predictor variable in both the count and zero inflation parts of the model. A linear model (package: MASS) was used to assess differences between treatment levels in log nibbling duration (s) (for those assays in which nibbling occurred). Variation in the occurrence of successful nibbling based on focal and opponent clerodanoid exposure (C+ or C−) was tested using a negative binomial model to account for overdispersion (glm.nb, package: MASS) with focal and opponent clerodanoid exposure (C+ or C−) and their interaction as predictor variables. A linear model was used to assess differences between treatments in log nibbling duration (s) where focal and opponent clerodanoid exposure (C+ or C−), and their interaction were the predictor variables.

### Pharmacophagy and conspecific transfer of clerodanoids

Binomial models (package: MASS, link=“logit”) were used to establish whether a) nibbling on *A. reptans* leaves and b) nibbling on conspecifics resulted in the acquisition of clerodanoids by *A. rosae*. The presence/absence of either of the two putative clerodanoid peaks from the chemical analyses was set as the response variable and either a) focal individual clerodanoid status (C−or C+) or b) non-focal individual clerodanoid status (C−or C+) and conspecific nibbling (Y/N) as the response variables. The relationship between nibbling duration and the sqrt of peak areas of each of the two compounds identified as putative clerodanoids was tested using a linear model.

## Results

### 1) Contest bioassays

The number of successful nibbling events was significantly higher in contest pairs where at least one individual was clerodanoid exposed (C+C+, C+C−) than in the control (C−C−) (Treatment;Log-likelihood: −215.02 on −4, X^2^= 18.951, p<0.001; Table 1), with the clerodanoid status of both the focal (X^2^_1,82_ = 4.058, p = 0.044) and opponent/non-focal individual (X^2^_1,82_ = 16.154, p < 0.001) having a significant, but non-interactive (X^2^_1,81_= 0.006, p = 0.94) effect on nibbling occurrence. Focal individuals were more likely to nibble on the opponent/non-focal individual when they were C−or the opponent/non-focal individual was C+ (Fig. 2). The nibbling duration was significantly affected by treatment (F_2,60_=5.801, p = 0.005), being longest in contest pairs in which one individual was clerodanoid exposed (‘C+C−’ −‘C+C+’, Z=2.658, p =0.021; ‘C+C−’ −‘C−C−’, Z = 2.800, p = 0.014), with the clerodanoid status of both the focal (F_1,43_= 4.918, p =0.032) and opponent/non-focal individual (F_1,43_ = 4.406, p = 0.042) having a significant, but non-interactive (F_1,42_ = 2.461, p = 0.124) effect on nibbling duration (Fig. 2).

**Table 1.** Results of zero-inflated negative binomial regression (negative binomial hurdle model) testing the effect of treatment on the occurrence of successful nibbling behaviour. Estimate is the estimated incident risk ratio (IRR) for the negative binomial model (‘count’) and the odds ratio (OR) for the logistic (zero inflation) model (‘zero’). Bootstrapped percentile adjusted lower (pLL) and upper (pUL) confidence intervals are also given.

| Parameter | Estimate | pLL | pUL |
| --- | --- | --- | --- |
| count (Intercept) | 4.15 | 2.91 | 5.35 |
| count C+vsC- | 2.06 | 1.42 | 3.11 |
| count C+vsC+ | 1.39 | 0.99 | 2.11 |
| zero (Intercept) | 1.00 | 0.38 | 2.60 |
| zeroC+vsC- | 3.40 | 1.08 | 12.10 |
| zero C+vsC+ | 9.50 | 2.18 | 4.54E+05 |

**Figure 2.**
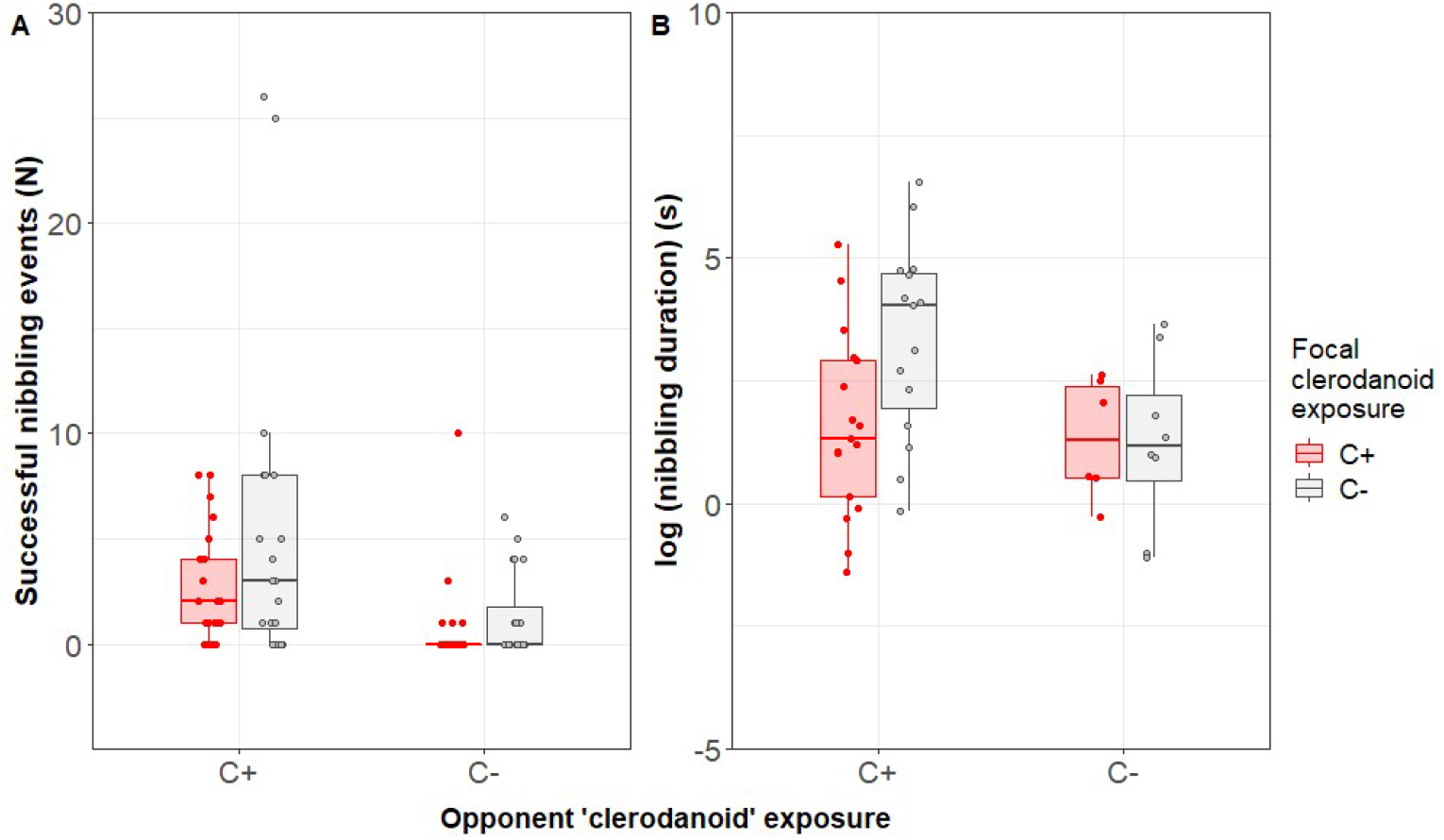
Effects of clerodanoid exposure of focal and opponent adults males of *A. rosae* (C+ = nibbled on *A. reptans* leaf, C−= not nibbled on *A. reptans* leaf) on the conspecific nibbling behaviour of the focal individual on another male [treatment (N):C+C+ (21), C−C+ (21),C+C−(23),C−C−(20)]. A) Number of successful nibbling events by the focal individual and B) total duration of all successful nibbling events. Boxes represent medians and interquartile ranges, whilst whiskers represent the smallest or largest values within 1.5 times the interquartile ranges either below the 25^th^ percentiles or above the 75^th^ percentiles, respectively.

### 2) Pharmacophagy and conspecific transfer of clerodanoids

Two putative clerodanoids were found in 75 % of the C+ animals but in none of the C− animals. The features found had 529.23 m/z and 527.21 m/z, which represent the [M+HCOOH-H]^-^ adducts of the candidate features 482.22 m/z (C_24_H_34_O_10_) and 484.23 m/z (C_24_H_36_O_10_). Only peaks consisting of at least four counts of the respective m/z were considered. The peaks of the candidate features, from now on referred to as 482 and 484, appeared in high intensities only in individuals that had direct or indirect access to *A. reptans* and the sum formulas suggested by the ‘SmartFormula’ function indicate that both features are clerodanoids. The candidates have a similar sum formula to ‘athaliadiol’ (C_24_H_34_O_9_) in the *Athalia rosae*-*Clerodendron trichotomum*-system (Nishida et al., 2004), which like the candidates was present in the insect extracts but not present in the plant material, indicating that they are equivalent metabolic products of plant compounds.

Nibbling on the leaves of *A. reptans* significantly influenced whether *A. rosae* individuals obtained the two clerodanoid compounds (X^2^_1,13_ = 11.193, p < 0.001). Nibbling of C−on C+ individuals significantly increased the likelihood of these individuals obtaining clerodanoids (AC+) compared to those who did not nibble on a C+ individual or that nibbled on a C− individual (clerodanoid status of opponent: X^2^_1,20_= 9.597, p = 0.002; nibbling occurrence: X^2^_1,20_ = 4.857, p = 0.027; Fig. 3). However, there was no significant effect of nibbling duration by C− individuals on the sqrt of the quantity of clerodanoids (= peak area) that they acquired for either of the two clerodanoids (compound 482: F_1,11_= 1.745, p= 0.213; compound 484: F_1,11_= 2.420, p=0.148).

**Figure 3.**
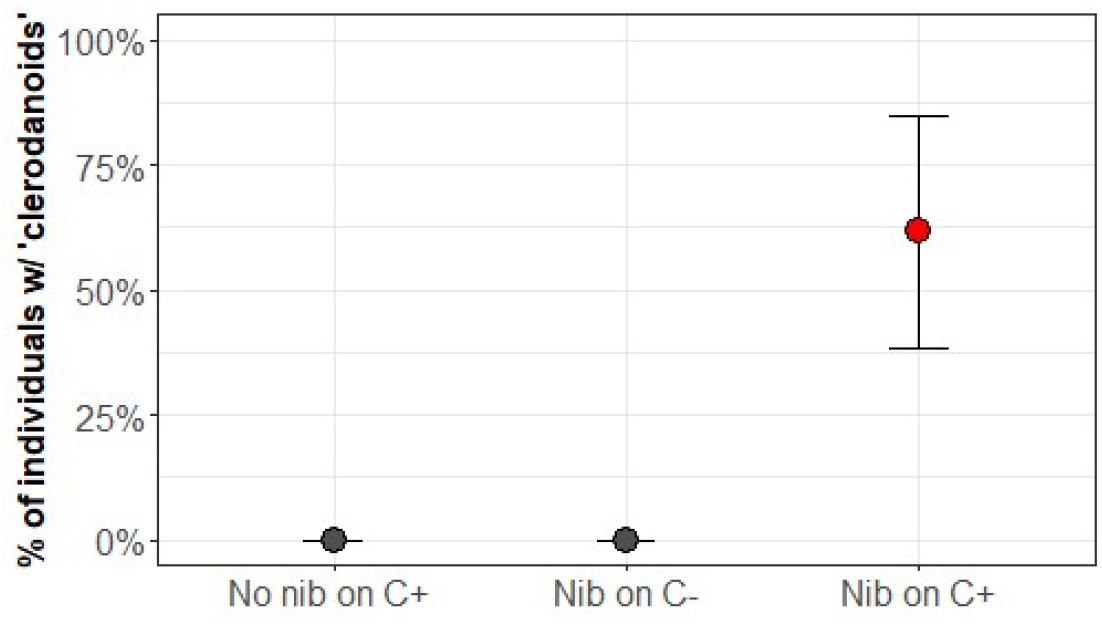
Percentage of C− males of *Athalia rosae*, in which clerodanoids (compounds 482 and 484) could be detected by chemical analysis after not nibbling on a C+ individual (N = 3), nibbling on a C− individual (N = 7), or nibbling on a C+ individual (N = 13), making them AC+.

### 3) Comparison of gene expression in C+, AC+ and C− individuals

More than 98% of reads were retained after cleaning. Prior to normalisation the average number of mapped reads per sample was 33,675,600, equating to an average of 91% reads per sample aligned to the predicted genes in the reference genome (range 85%-93%). Access to clerodanoids via conspecific nibbling (AC+) had the strongest effect on gene expression (Figs. 4 & 5). Between males that had nibbled on conspecifics (AC+) and those that had either nibbled on *A. reptans* leaves (C+) or had no access to a source of clerodanoids (C−) there were respectively 318 (AC+ vs C+) and 766 (AC+ vs C−) significantly differentially expressed genes (LFC > 0, adjusted p < 0.05). There were considerably fewer (50) differentially expressed genes between C+ and C− individuals (C + vs C−). Few differentially expressed genes were found that belonged to previously identified canonical sequestration and detoxification pathways, namely P450s, GSTs, UDPs, or ABC transporters (as identified by gene name). For example, between C+ and C− males, in which one might have expected such genes to be expressed, only one gene, from the ABC transporter category (ABCD1/LOC105692744), was significantly upregulated in C+ individuals, but even then the transporter is linked predominantly to long chain fatty acid transport. In comparison, many more P450s, GSTs, UDPs, or ABC transporters genes were significantly upregulated in AC+ individuals than in both C+ and C− individuals (S4). In particular two cytochrome P450s were upregulated alongside a raft of cytochrome genes linked to mitochondrial function (S4). In terms of those DE genes shared between AC+ vs C− and AC+ vs C+, which may give a strong indication of genes linked to conspecific nibbling behaviour (166 DE genes, Fig. 4), the *takeout* gene (to/CG11853) and *flightin* gene (fln/CG7445) were strongly upregulated in AC+ (S5). The former has been linked to both adult feeding and courtship behaviour in *Drosophila melanogaster* (Dauwalder et al., 2002; Sarov-Blat et al., 2000) and the latter is thought to be involved in the regulation of flight muscle contraction (Henkin et al., 2004). Interestingly one of the genes shared by AC+ vs C− and C+ vs C− differential expression, which may give a strong indication of genes linked to pharmacophagy (24 DE genes, Fig. 4), was *tdc* the gene encoding L-tyrosine decarboxylase, which has been linked to stress responses, acid resistance and drug metabolism (Perez et al., 2015; van Kessel et al., 2019), but again no canonical detoxification genes were shared (S5).

**Figure 4.**
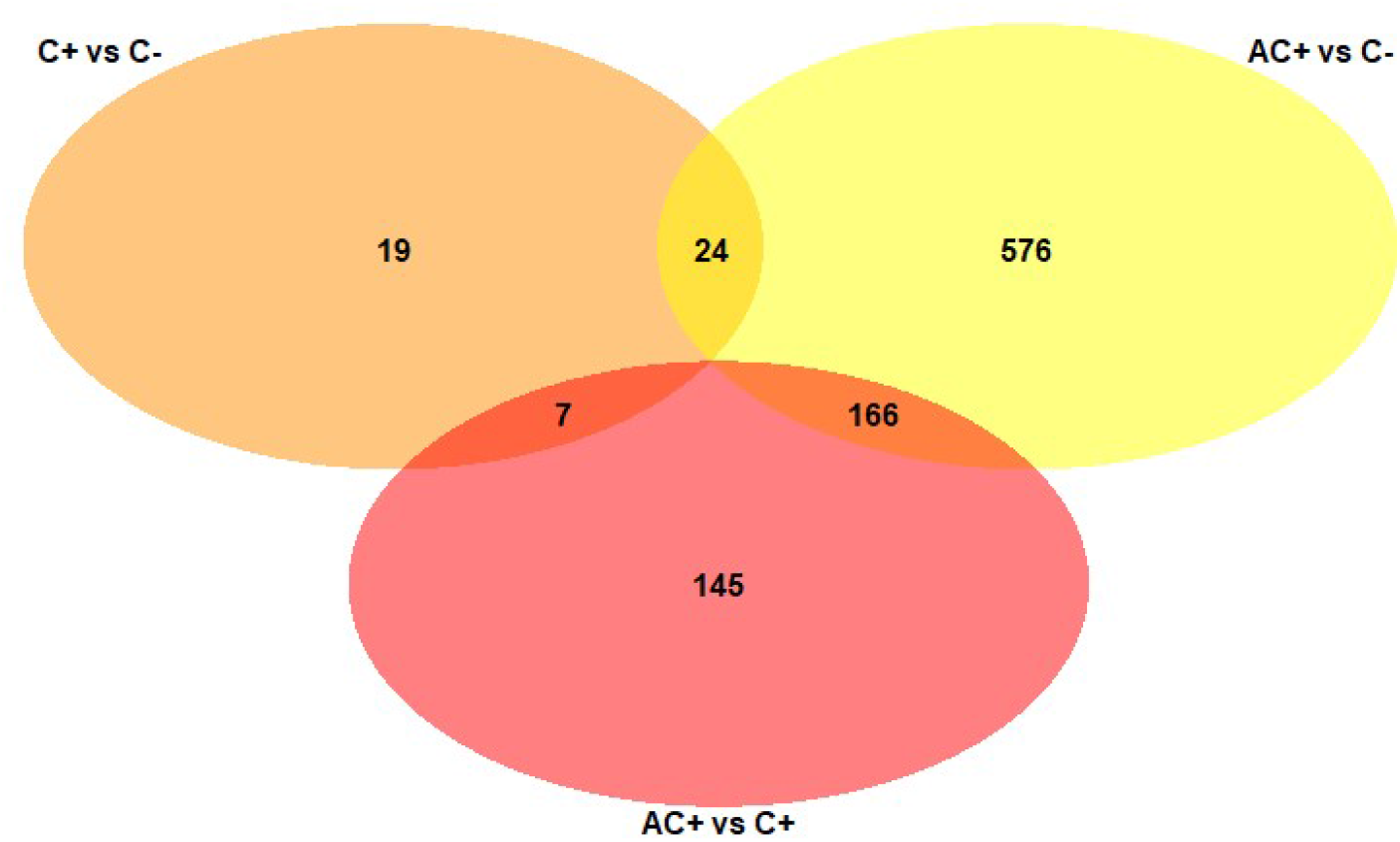
Venn diagram illustrating the number of unique and shared differentially expressed genes resulting from each of the three pairwise comparisons (AC+ vs C+, red; AC+ vs C−, yellow; C+ vs C−, orange), in which male *Athalia rosae* had access to clerodanoids via conspecific nibbling (AC+), nibbling on *A. reptans* leaves (C+), or had no access to a source of clerodanoids (C−).

**Figure 5.**
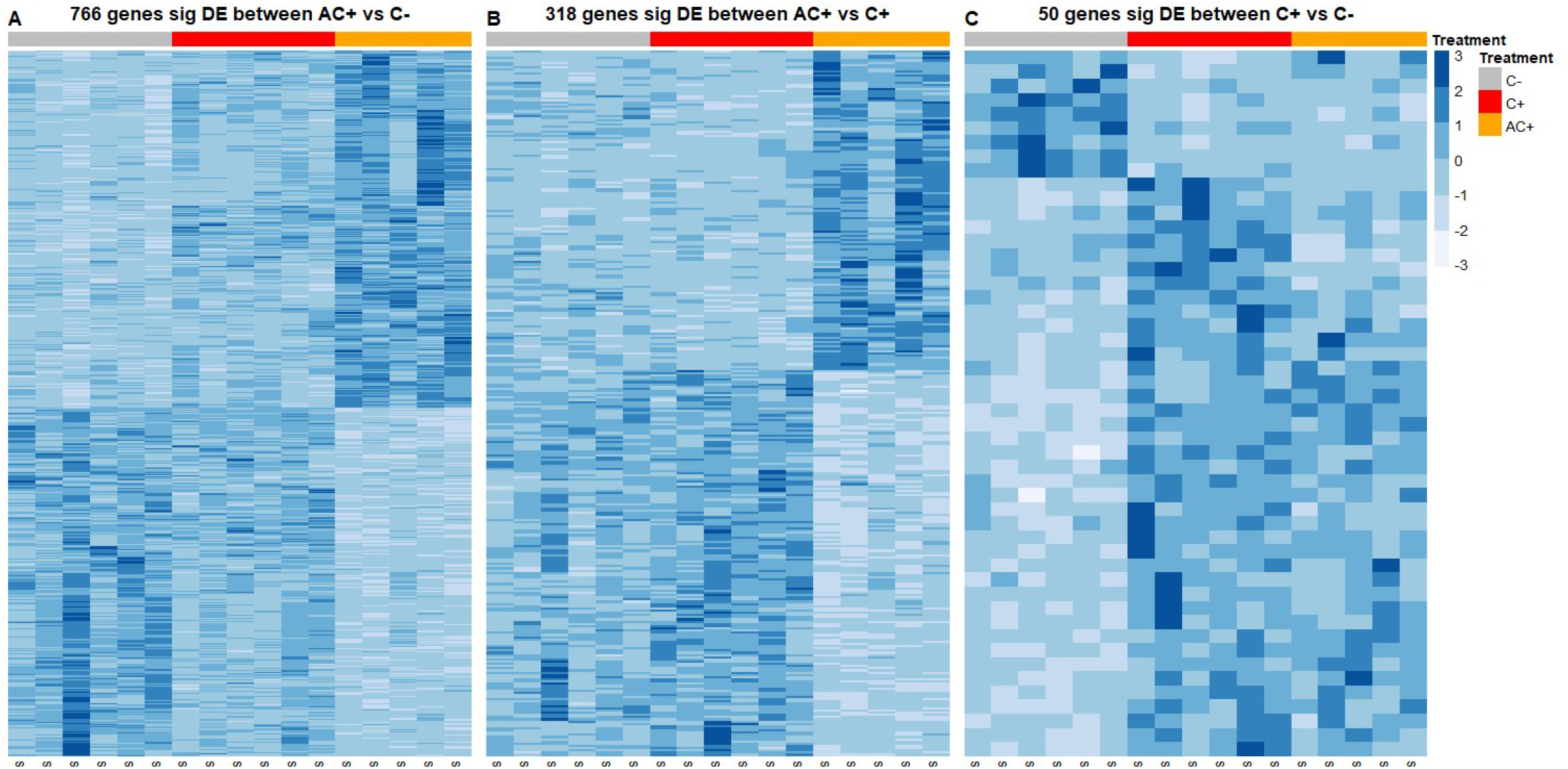
Heatmap showing the expression of genes (scaled normalised counts) across all samples for those **A**) 766 genes that were significantly (sig) differentially expressed (DE) between (AC+) and (C−), **B**) 318 genes that were significantly DE between (AC+) and (C+), **C**) 50 genes that were significantly DE between (C+) and (C−). AC+ = adult male *Athalia* rosae that had access to clerodanoids via nibbling on conspecifics, C+ = *A. rosae* that had access to clerodanoids via nibbling on *Ajuga* reptans leaves, and C−= those that had no access to a source of clerodanoids.

Whilst the majority of reads mapped to the reference genome, 247,997,896 reads did not map; we assembled these into 334,717 transcripts using Trinity. We further explored those with read coverage >10,000 reads (791 transcripts). Of those, 688 were putatively annotated against the NCBI nt database, and the majority were mitochondrial. DE analysis of unmapped reads showed that a low number of genes were differentially expressed between the treatment pairs 136 for AC+ vs C−, 60 for AC+ vs C+, and 10 for C+ vs C−. Of these significantly DE unmapped genes very few had putative annotations (9, 7, and 2 genes respectively) and in turn few of these had identities linked to *A. rosae* (4, 2, and 0 genes respectively).

Gene set enrichment analysis using KEGG annotations revealed that AC+ showed the highest number of non-redundant differentially expressed gene sets or pathways compared to both of the other treatments and that many of these pathways were shared between the comparisons of AC+ with the two other treatments (Table 2; see S6 for full output). The pathways upregulated in AC+ fall roughly into three categories: energy and carbohydrate metabolism (‘oxidative phosphorylation’, ‘carbon metabolism’, ‘pentose phosphate pathway’, and ‘fructose and mannose metabolism’), endocrine systems (‘glucagon signalling pathway’ and ‘thyroid hormone signalling pathway’), and circulatory systems (‘cardiac muscle contraction’ and ‘adrenergic signalling in cardiomyocytes’). The cGMP-PKG signalling pathway was also significantly upregulated in AC+ individuals compared to both the control (C−) and C+ and plays a role in a number of processes including muscle contraction and memory formation in insects (Matsumoto et al., 2018). Furthermore, the focal adhesion pathway, the turnover of which increases with stress (Aedo et al., 2019), was significantly upregulated in AC+ individuals compared to the control (C−). Finally, the ‘ribosome’ pathway was significantly downregulated in AC+ individuals in comparison to the two other treatments and interestingly it was also downregulated in C+ compared to the control (C−).

**Table 2.**
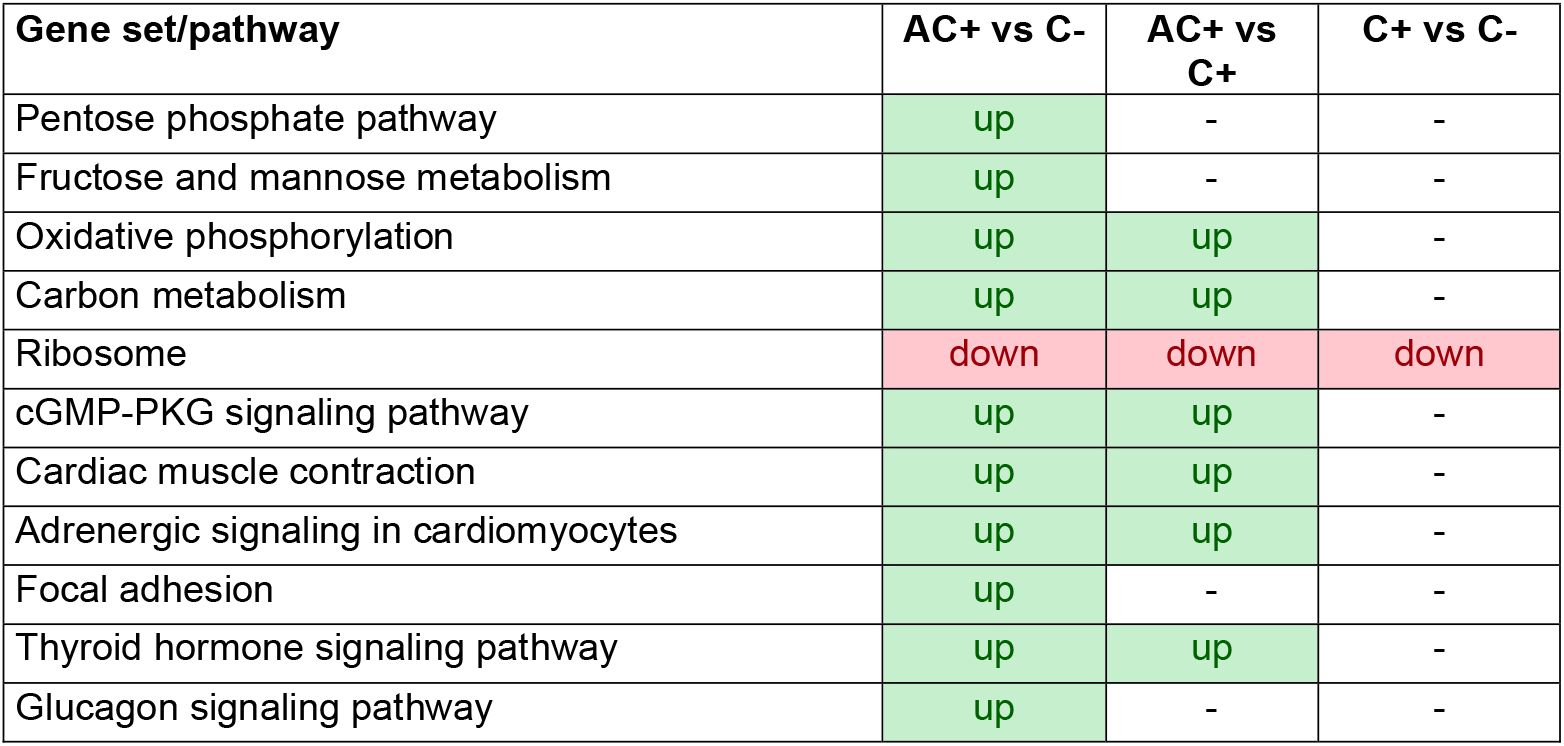
Significantly differentially expressed KEGG gene sets/pathways between adult males of *A. rosae* of the three clerodanoid treatment levels. Left hand side is the ‘treatment’ and right hand side the ‘control’ in each comparison, meaning that pathways are up-or downregulated in ‘treatment’ relative to ‘control’. AC+ = adult male *Athalia rosae* that had access to clerodanoids via nibbling on conspecifics, C+ = *A. rosae* that had access to clerodanoids via nibbling on *Ajuga reptans* leaves, and C−= those that had no access to a source of clerodanoids

| Gene set/pathway | AC+ vs C- | AC+ vs C+ | C+ vs C- |
| --- | --- | --- | --- |
| Pentose phosphate pathway | up | - | - |
| Fructose and mannose metabolism | up | - | - |
| Oxidative phosphorylation | up | up | - |
| Carbon metabolism | up | up | - |
| Ribosome | down | down | down |
| cGMP-PKG signaling pathway | up | up | - |
| Cardiac muscle contraction | up | up | - |
| Adrenergic signaling in cardiomyocytes | up | up | - |
| Focal adhesion | up | - | - |
| Thyroid hormone signaling pathway | up | up | - |
| Glucagon signaling pathway | up | - | - |

## Discussion

Here we investigated a novel form of pharmacophagy in adults of the sawfly *A. rosae* and show that individuals are able to acquire compounds known to influence defence and mating behaviour (Amano et al., 1999; Nishida and Fukami, 1990), not only from plants (*A. reptans* leaves) but also from the bodies of conspecifics who have previously fed on those plants. In contrast to plant-derived nuptial gifts that are usually given from males to females (Zagrobelny et al. 2007, Wink 2019) the transfer of plant-derived chemicals described here is occurring within and between both sexes in this sawfly species predominantly in the context of dyadic contest behaviour (this study and Paul et al *in prep*). Thus, acquisition of plant compounds affects the metabotype of the individuals, which has consequences for social interactions with conspecifics.

The contest pairs in this experiment were matched for size, as a proxy for RHP (McCann, 1981), and as such the main determinant of agonistic interactions between individuals was most probably the value of the resource to each individual (RV) and therefore the motivation of each male to fight (Arnott and Elwood, 2008; Stockermans and Hardy, 2013). As predicted, the occurrence and duration of nibbling by focal individuals increased when they themselves had not previously collected clerodanoids (C−). This indicates that C− individuals had a higher motivation than C+ individuals to attack and agonistically nibble on C+ opponents. Furthermore, although C− focal individuals were more likely to successfully nibble an opponent, C+ focal individuals still engaged in the successful nibbling of opponents when those opponents were also C+. Therefore, contrary to predictions, it appears that focal individuals will nibble on C+ opponents both to gain (focal = C−) and to potentially increase (focal = C+) the quantity of clerodanoids that they possess.

Fighting between adult *A. rosae* over clerodanoids is likely to carry many of the costs associated with contest interactions, including energy expenditure (Hack, 1997), accruing of injuries (Lane and Briffa, 2017), and a potential increase in predation risk (Baker et al., 1999) caused by reduced vigilance during a contest (Jakobsson et al., 1995). That focal C+ individuals still engaged in agonistic behaviour and aggressively nibbled on C+ opponents despite these costs indicates the high value of accruing more clerodanoids. It also suggests that the beneficial effects of clerodanoids might be concentration-dependent, a common attribute of specialised metabolites in insect defence (Burdfield-Steel et al., 2019), immunity (Tan et al., 2019), and mate attraction (Iyengar et al., 2001; Kelly et al., 2012). Future work may therefore assess to what extent clerodanoid concentrations and the overall metabotype of *A. rosae* changes in dependence of contact intensity and how this might influence social interactions, both fighting and mating.

It has previously been demonstrated in *A. rosae ruficornis* that adults that nibbled on *A. reptans* leaves acquired a clerodanoid-like compound (athaliadiol, C24H34O9) that differed from the clerodanoids taken up from leaves, and which was not present in those individuals that had not nibbled on leaves (Nishida et al., 2004). Our data likewise indicate that plant compounds are further metabolised by the sawflies, as via our chemical analyses we found two putative clerodanoids in C+ animals but not in the leaves. Metabolism of plant specialised metabolites occurs in many insects after uptake to lower the toxicity of these compounds and increase their solubility (Erb and Robert, 2015). However, no significant upregulation of genes belonging to canonical detoxification and sequestration pathways could be found in C+ *A*.*rosae* individuals. In comparison, in AC+ individuals two cytochrome P450 genes were upregulated compared to the control (C−). Cytochrome P450 genes often play a critical role in phase 1 of the metabolism and sequestration of plant specialised metabolites through the oxidation, reduction, or hydrolysis of compounds (Birnbaum and Abbot, 2020). One of the two genes upregulated in AC+ individuals, *Cyp4c3*, has been linked to xenobiotic resistance in *Drosophila* (Scanlan et al., 2020) and many genes in the insect CYP4 family play a role in detoxification in insects (Scott, 1999; Vogel et al., 2014). Despite this, no other genes or pathways directly linked to detoxification or sequestration were upregulated in AC+. These findings may in part be related to the timing of the experiment. Upregulation of such genes may be rapidly induced when the uptake of specialised compounds is initiated and may level down shortly after and therefore fall outside of our sampling period.

An alternative explanation is that the clerodanoid compounds, although shown to be distasteful (Nishida and Fukami, 1990), might not actually be toxic and therefore may not require the same metabolism as highly toxic compounds. Both distasteful and toxic compounds play a role in insect defence mechanisms (Marples et al., 2018), and the ingestion of either may be associated with different metabolic signatures. Finally, it is unclear at this point to what extent clerodanoids are indeed ingested by *A. rosae*, in contrast to the larvae of this species, which sequester glucosinolates from the ingested plant material of their Brassicaceae hosts in their haemolymph (Müller et al., 2001). In the adults, the clerodanoids must be present on the exterior of *A. rosae*, as conspecifics can acquire them through nibbling on the exoskeleton of C+ individuals. Thus, clerodanoids may simply be collected using the mouthparts, metabolised by salivary glands (as seen in species of Hemiptera, e.g. Zhu et al., 2016) or alternatively break down non-enzymatically once liberated from the cellular compartments in *A. reptans* leaves. These modified clerodanoids may then be redistributed over the body via grooming behaviour (SCP pers. obs.). Further work on the fate of clerodanoids at different time points after leaf pharmacophagy commences and in different parts of the *A. rosae* body, as done for glucosinolates in *A*.*rosae* larvae (Abdalsamee et al., 2014), is therefore needed to help to clarify the mechanisms at play.

Taken together, the evidence indicates that the costs of clerodanoid uptake in *A. rosae* through leaf nibbling are low, at least in terms of metabolic costs. This fits the wider picture of research on sequestration costs, which apart from a few exceptions (Birnbaum et al., 2017; Camara, 1997; Cogni et al., 2012) seem to be negligible (Zvereva and Kozlov, 2016). In comparison the metabolic costs of acquiring clerodanoids from conspecifics (opposed to leaves) appears to be much higher, as illustrated by the upregulation of gene pathways related to carbon metabolism and ATP production in AC+ individuals. This is most likely related to the fact that acquisition from conspecifics involves fighting and fighting is known to be energetically costly (Briffa and Sneddon, 2007; Hack, 1997). For example, in hermit crabs glycogen reserves are mobilized to an increasing degree during a fight as fight intensity escalates (Briffa and Elwood, 2004). Although it is important to note that we have not translated these metabolic costs into fitness costs, the fact that individuals are willing to forgo the cost of fighting to gain clerodanoids, even when they already have some, indicates that such costs are heavily outweighed by the fitness benefits of gaining clerodanoids. These benefits may not just be limited to their role in mate choice (Amano et al., 1999) and predator deterrence (Nishida and Fukami, 1990), but also encompass antimicrobial defence. Clerodanoids are known to act in an antimicrobial manner (Bozov et al., 2015) and consequently may positively influence fitness by being used as self-medication, as known from other sequestered chemicals (Abbott, 2014; Tan et al., 2019). A more holistic view of the effects of clerodanoids on fitness would therefore help us to understand what might be driving such apparently costly nibbling behaviour.

In addition to the upregulation of gene pathways related to the increased metabolic demands being placed on the body during a fight in AC+ individuals, expression of ribosomal proteins was downregulated between AC+ and the other two treatments, and C+ and the control treatment (AC+ < C+ < C−). Ribosomal proteins (RPs) play a key role in ribosome assembly, stability, and translation, and as a consequence the genes and gene pathways responsible for RPs are highly conserved and their expression was assumed to be relatively stable. However, it is becoming increasingly clear that this is not the case and that the expression of RPs can change in response to a number of different factors, e.g. heat stress (Paraskevopoulou et al., 2020) or the ingestion of ribosome-inactivating proteins (RIPs) contained in plant material (Celorio-Mancera et al., 2013; Stirpe, 2013). RPs also play numerous roles outside of ribosomal biogenesis (Zhou et al., 2015). For example, RP expression might be lower in AC+ and C+ individuals because of their role in immune response. Multiple RPs have been shown to play a role in innate immune responses, e.g. *RPL13A* (Mukhopadhyay et al., 2008; Mukhopadhyay et al., 2009), and there is evidence in monarch butterflies (*Danaus plexippus*) that the acquisition of plant specialised metabolites by pharmacophagy aids fighting off parasites whilst resulting in the downregulation of several genes that play a role in immunity (Tan et al., 2019). The acquisition of clerodanoids may therefore work in a similar way, increasing general immunity due to their previously mentioned antimicrobial effects (Bozov et al., 2015) and leading to the downregulation of genes involved in innate immunity including ribosomal proteins. Furthermore, in AC+ individuals the greater downregulation of RPs (when compared to both C+ and C−) could additionally reflect the sensitivity of ribosome biogenesis to the energy status of the cell (Zhou et al., 2015), with nutrient deprivation previously being demonstrated to result in the degradation of RPs (Kraft et al., 2008). Although AC+ individuals were not nutrient-deprived, fighting is energetically expensive and the downregulation of RPs may well result as a mechanism to conserve ATP.

In conclusion here we show that pharmacophagous behaviour by insects occurs on both plants and conspecifics and that the acquisition of specialised plant metabolites and consequent changes in an individual’s metabotype has ramifications for social interactions and therefore potentially their social niche. Gene expression in *A. rosae* indicates that in this species metabolic costs are low for clerodanoid sequestration from plants but high for individuals collecting clerodanoids from conspecifics by agonistically nibbling on their exterior. Conspecifics were willing to bear these nibbling costs even if they had already acquired some clerodanoids themselves. Further work is needed in order to fully elucidate the metabolic fate of clerodanoids in *A. rosae* and how the putative costs and benefits of gaining and fighting for clerodanoids identified here translate into fitness effects.

## Author contributions

Conceptualization: SCP, CM; Methodology: SCP, ABD, LJT; Investigation: SCP, LJT, JF; Resources: CM, ABD; Formal analysis and validation: SCP, ABD, LJT; Data Curation Management: SCP, LJM; SCP wrote the manuscript and CM, ABD, and LJT contributed substantially to the final version.

## Acknowledgements

Many thanks to the University of Potsdam AG Genetics and AG Evolutionary Adaptive Genomics for use of computational resources and to Dr Tobias Busche and Katrin Lehmann for assistance with the RNA extraction.

## Data availability

Data and code for behavioural analysis (https://doi.org/10.5281/zenodo.4523139) and for gene expression analysis (https://doi.org/10.5281/zenodo.4523881) accessible via ZENODO. Raw Reads: BioProject ID = PRJNA700827. BioSample accession numbers = SAMN17839831-48 (http://www.ncbi.nlm.nih.gov/bioproject/700827)

