## Supplementary material for "Consequences of pharmacophagous uptake from plants and conspecifics in a sawfly elucidated using chemical and molecular techniques": S1

**Supplement 1 –** UHPLC-QTOF-MS/MS settings

From the extracts, 8 µl were applied to the column and separated at 45°C on a gradient from 0.1% formic acid (p.a., eluent additive for LC-MS, ~98 %, Sigma-Aldrich; in millipore water) to 0.1 % formic acid in acetonitrile (LC-MS grade, Fisher Scientific, Loughborough, UK; eluent B) at a flow rate of 0.5 ml min^-1^. The proportion of eluent B started at 2% and increased to 30% B within 20 min, and further to 75% B within 9 min. Electrospray ionization was done with the source settings: end plate offset: 500 V, capillary voltage: 3000 V, nebuliser (N_2_) pressure: 3 bar, dry gas (N_2_) flow and temperature: 12 l min^-1^ at 275 °C. Line spectra (50-1300 m/z) were acquired at a spectra rate of 1 Hz. A Na(HCOO)-based calibration solution was introduced to the source at the end of each sample for mass axis recalibration. The quadrupole settings were: ion energy: 4 eV, low mass: 90 m/z, and the collision cell settings were: collision energy: 7 eV, transfer time: 100 µs, pre-pulse storage: 5 µs.
